# Endogenous insulin contributes to pancreatic cancer development

**DOI:** 10.1101/530097

**Authors:** Anni M.Y. Zhang, Jamie Magrill, Twan J.J. de Winter, Xiaoke Hu, Søs Skovsø, David F. Schaeffer, Janel L. Kopp, James D. Johnson

## Abstract

Obesity and early-stage type 2 diabetes (T2D) increase the risk for many cancers, including pancreatic ductal adenocarcinoma (PDAC). The mechanisms linking obesity and T2D to cancer have not been established, preventing targeted interventions. Arguments have been made that hyperinsulinemia, hyperglycemia, or inflammation could drive cancer initiation and/or progression^1^. Hyperinsulinemia is a cardinal feature of obesity and T2D, and is independently associated with PDAC incidence and mortality^2–4^, even in non-obese people^5^. Despite ample human epidemiological evidence linking hyperinsulinemia to PDAC, there is no direct *in vivo* evidence of a causal role for endogenous insulin in cancer in any system. Using mice with reduced insulin gene dosage^6, 7^, we show here that a modest reduction in endogenous insulin production leads to a ~50% reduction in pancreatic intraepithelial neoplasia (PanIN) pre-cancerous lesions in high fat diet-fed mice expressing the *Kras*^G12D^ oncogene^8^. The significant reduction in PanIN lesions occurred in the absence of changes in fasting glucose. Reduced insulin also led to a ~50% reduction in pancreatic fibrosis, suggesting that endogenous insulin drives PanIN development, in part, via its pro-fibrotic effects on the stroma surrounding acinar cells and PanIN. Collectively, our data indicate that endogenous insulin hypersecretion contributes causally to pancreatic cancer development. This suggests a modest reduction in fasting insulin via lifestyle interventions or therapeutics may be useful in cancer prevention.

---

The incidence of many cancers, including pancreatic cancer, continues to rise alongside the global obesity and T2D epidemics^9^. There is also increasing popular interest in the role of diet and lifestyle in cancer risk and prevention strategies. Low carbohydrate diets have been proposed for prevention or as an adjunct to cancer therapy based on evidence that many tumours selectively use glucose as fuel^10^ or based on what has come to be known as the insulin-cancer hypothesis^11, 12^. Insulin has mitogenic and anti-apoptotic effects on pancreatic cell cultures in a dosage-dependent manner^13^ and is known to activate signalling pathways implicated in tumourigenesis^13, 14^. However, despite the abundance of clinical evidence of association between insulin and various cancer types^2–4, 15^, the insulin-cancer hypothesis has never been formally tested. Testing the causality of hyperinsulinemia in cancer requires a controlled *in vivo* system susceptible to tumor development wherein circulating insulin can be controlled, ideally independently of glucose. Mutations in the *KRAS* proto-oncogene are found in 90% of human pancreatic cancers and, when present in pancreatic exocrine cells, induces PanIN lesions and PDAC reliably in mice^8^. PanIN are non-invasive preneoplastic mucinous lesions with ductal morphology that are considered precursors to PDAC^16^. Therefore, to test the hypothesis that a diet-induced increase in circulating insulin plays a causal role in the initiation of PanIN and PDAC, we expressed oncogenic Kras in murine acinar cells (*Ptf1a*^CreER^;LSL-*Kras*^G12D^ mouse model^8, 17^) to promote induction of PanIN in the presence or absence of reduced insulin production^7, 18^.

We controlled insulin production by generating control mice with a normal complement of two wildtype *Ins1* alleles (*Ins1*^+/+^) and experimental mice with only a single functional *Ins1* allele (*Ins1*^+/−^) from the same sets of parents (Fig. 1a). In our experiments, all mice were maintained on an *Ins2* null (*Ins2*^−/−^) background to prevent insulin gene compensation from the secondary copy of the *Insulin* gene present in mice^19^. We have previously found that these genetic manipulations of the *Ins1* and *Ins2* loci result in a sustained reduction in fasting insulin that is modest enough not to chronically impair glucose homeostasis^7, 18^. Mice were weaned onto a high-fat diet (HFD) (Fig. 1b) known to cause hyperinsulinemia, accelerate PanINs formation, and increase tumour incidence in *Kras*^G12D^ expressing PDAC mice^1, 20^. Control and experimental mice were injected with tamoxifen around 7 weeks of age to induce *Kras*^G12D^ expression and maintained on HFD for 1 year (Fig. 1b).

**Fig. 1.**
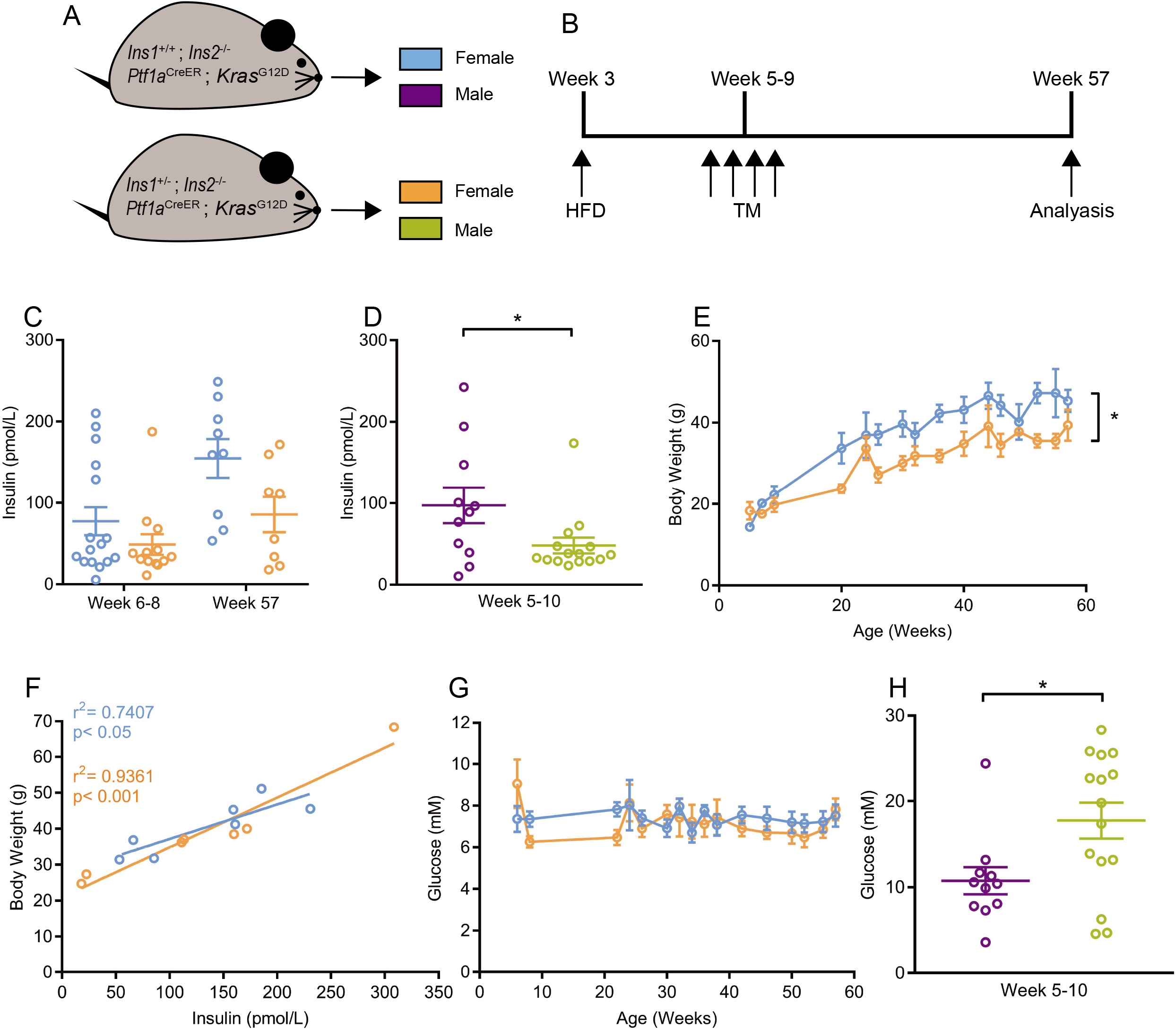
Fasting insulin, glucose and body weight in mice with reduced insulin gene dosage. **a**, Schematic describing the mouse model used to test whether hyperinsulinemia contributes causaly to high-fat diet (HFD)-promotion of PDAC progression. **b**, Schematic describing the experimental design, briefly 3-week-old control and experimental mice were weaned to HFD and then injected on 4 consecutive days with tamoxifen (TM) beginning at 5-9 weeks. Mice were euthanized at 57 weeks of age for analysis. **c-d**, Fasting circulating insulin levels of female (c) (n= 16 and 9 for *Ptf1a*^CreER^;LSL-*Kras*^G12D^;*Ins1*^+/+^;*Ins2*^−/−^ mice at week 6-9 and week 57, respectively. n= 13 and 8 for *Ptf1a*^CreER^;LSL-*Kras*^G12D^;*Ins1*^+/−^;*Ins2*^−/−^ mice at week 6-9 and week 57, respectively.) and male (d) (n= 11 and 15 for *Ptf1a*^CreER^;LSL-*Kras*^G12D^;*Ins1*^+/+^;*Ins2*^−/−^ mice and *Ptf1a*^CreER^;LSL-*Kras*^G12D^;*Ins1*^+/−^;*Ins2*^−/−^ mice, respectively) mice of the indicated genotypes and at the indicated time points. P vaules comparing *Ptf1a*^CreER^;LSL-*Kras*^G12D^;*Ins1*^+/+^;*Ins2*^−/−^ to *Ptf1a*^CreER^;LSL-*Kras*^G12D^;*Ins1*^+/−^;*Ins2*^−/−^ mice were 0.3744 by Mann-Whitney test, 0.0509 by two-tailed unpaired t-test, and 0.0403 (*) by Mann-Whitney test for female at week 6-8, female at week 57, and male at week 5-10, respectively. e, Body weights were measured over 1 year for female mice (n= 16 and 13 for *Ptf1a*^CreER^;LSL-*Kras*^G12D^;*Ins1*^+/+^;*Ins2*^−/−^ mice and *Ptf1a*^CreER^;LSL-*Kras*^G12D^;*Ins1*^+/−^;*Ins2*^−/−^ mice, respectively.). P<0.0001 (*) by two-way ANOVA. **f**, Relationship between body weight and insulin at 1 year of age (n= 7). **g**, Periodic measurements of 4-hour fasted blood gucose over 1 year for female mice (n= 16 and 13 for *Ptf1a*^CreER^;LSL-*Kras*^G12D^;*Ins1*^+/+^;*Ins2*^−/−^ mice and *Ptf1a*^CreER^;LSL-*Kras*^G12D^;*Ins1*^+/−^;*Ins2*^−/−^ mice, respectively.). **h**, The 4-hour fasted blood glucose at week 5-10 for male mice (For *Ptf1a*^CreER^;LSL-*Kras*^G12D^;*Ins1*^+/+^;*Ins2*^−/−^ mice, n= 11 and for *Ptf1a*^CreER^;LSL-*Kras*^G12D^;*Ins1*^+/−^;*Ins2*^−/−^ mice, n= 15.) P vaules comparing *Ptf1a*^CreER^;LSL-*Kras*^G12D^;*Ins1*^+/+^;*Ins2*^−/−^ to *Ptf1a*^CreER^;LSL-*Kras*^G12D^;*Ins1*^+/−^;*Ins2*^−/−^ mice by Mann-Whitney test was 0.0191 (*).). Values are shown as mean ± SEM.

First, we characterized glucose homeostasis in *Ins1*^+/−^;*Ins2*^−/−^ mice and *Ins1*^+/+^;*Ins2*^−/−^ mice on the *Ptf1a*^CreER^;LSL-*Kras*^G12D^ genetic background (Fig. 1b). In both female and male mice, there was an expected reduction in fasting insulin levels in *Ptf1a*^CreER^;LSL-*Kras*^G12D^;*Ins1*^+/−^;*Ins2*^−/−^ mice with reduced insulin gene dosage compared to *Ptf1a*^CreER^;LSL-*Kras*^G12D^;*Ins1*^+/+^;*Ins2*^−/−^ controls (Fig. 1c-d). Consistent with our previous study on a different background^7^, we found reducing *Ins1* gene dosage in females caused a reduction in body weight and body weight correlated with circulating insulin levels (Fig. 1e-f). Importantly, this partial reduction in insulin produced in *Ptf1a*^CreER^;LSL-*Kras*^G12D^;*Ins1*^+/−^;*Ins2*^−/−^ mice did not affect fasting glucose levels in female mice over the one year study (Fig. 1g). In male mice from the same colony, we observed significantly impaired fasting glucose, and in some cases frank diabetes (Fig. 1h), consistent with the established sex differences in insulin sensitivity. The high incidence of diabetes in males prevented us from discriminating the specific effects of insulin in males over the planned study period and led us to focus primarily on female mice. Thus, comparing female *Ptf1a*^CreER^;LSL-*Kras*^G12D^;*Ins1*^+/−^;*Ins2*^−/−^ experimental mice to female *Ptf1a*^CreER^;LSL-*Kras*^G12D^;*Ins1*^+/+^;*Ins2*^−/−^ controls offered us the unique opportunity to test the role of insulin in pancreatic cancer initiation, in the absence of changes in fasting glucose.

To test our primary hypothesis that decreasing endogenous insulin production would affect the initiation of *Kras*^G12D^-driven PDAC, we measured the percent of total pancreatic area occupied by PanIN and tumour in hematoxylin and eosin (H&E) stained pancreatic sections. At one-year, abundant ductal lesions with histologic and molecular characteristics of low-grade PanINs (Fig. 2a), including highly acidic mucin content indicated by Alcian blue staining (Fig. 2b), were found in both groups. Consistent with previous studies^8^, *Kras*^G12D^ expression in the pancreas also resulted in rare PDAC, which expressed ductal marker Cytokeratin 19 (CK19) (Fig. 2c-d). However, the incidence of PDAC was similarly rare in both genotypes. Therefore, we focused on the effect of insulin on the initiation of PanIN lesions. We reasoned that if hyperinsulinemia played a causal role in cancer formation in the context of diet-induced obesity then mice with reduced insulin would have a smaller combined area occupied by PanIN lesions and tumours. Indeed, experimental *Ptf1a*^CreER^;LSL-*Kras*^G12D^;*Ins1*^+/−^;*Ins2*^−/−^ mice had approximately half the area covered by PanIN or tumour compared with *Ptf1a*^CreER^;LSL-*Kras*^G12D^;*Ins1*^+/+^;*Ins2*^−/−^ control mice (12.76 ± 3.46% vs 25.40 ± 3.82%; p=0.0306) (Fig. 2c). This indicates the reduction in endogenous insulin production significantly reduced PanIN initiation. Furthermore, PanIN plus tumour area in both groups of mice tended to correlate with their fasting insulin levels at 1 year of age (Fig. 2e). PanIN area also correlated with body weight at 1 year of age in control *Ptf1a*^CreER^;LSL-*Kras*^G12D^;*Ins1*^+/+^;*Ins2*^−/−^ mice, but not *Ptf1a*^CreER^;LSL-*Kras*^G12D^;*Ins1*^+/−^;*Ins2*^−/−^ mice (Fig. 2f). For both genotypes, the percent PanIN area did not correlate with 4-h fasted glucose (Fig. 2g), which was stable throughout the study period (Fig. 1g). Therefore, this further argues against a prominent role for glycemia in PanIN formation. Although we were unable to follow them for 1 year, qualitatively similar data were observed in male mice (Fig. 2h-i). The percent PanIN area tended to correlate with the ages of the mice (Fig 2j). Together, our data implicate endogenous increases in insulin, but not glucose, in HFD-mediated promotion of PanIN development and potentially pancreatic cancer.

**Fig. 2.**
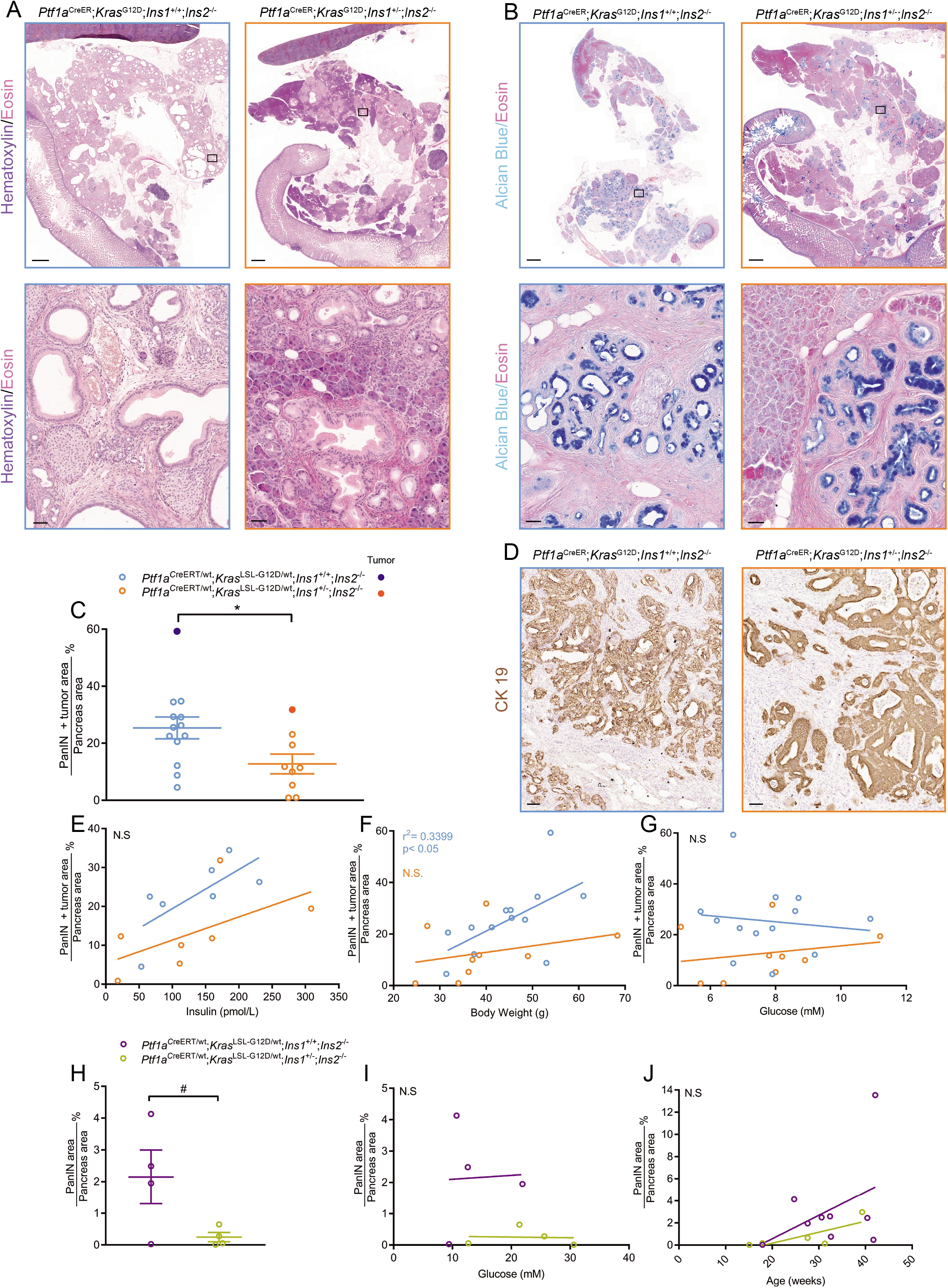
Reduced PanIN/tumour area in mice with reduced insulin production. **a**, Representative whole-section (top) and high-magnification (bottom) images of *Ptf1a*^CreER^;LSL-*Kras*^G12D^;*Ins1*^+/+^;*Ins2*^−/−^ and *Ptf1a*^CreER^;LSL-*Kras*^G12D^;*Ins1*^+/−^;*Ins2*^−/−^ mice pancreata stained with H&E. **b**, Representative whole-section (top) and high-magnification (bottom) images of *Ptf1a*^CreER^;LSL-*Kras*^G12D^;*Ins1*^+/+^;*Ins2*^−/−^ and *Ptf1a*^CreER^;LSL-*Kras*^G12D^;*Ins1*^+/−^;*Ins2*^−/−^ mice pancreata stained with Alcian blue and eosin. **c**, Quantification of percent of total pancreatic area occupied by PanIN and tumour in female mice of each genotype (n= 13 and 9 for *Ptf1a*^CreER^;LSL-*Kras*^G12D^;*Ins1*^+/+^;*Ins2*^−/−^ mice and *Ptf1a*^CreER^;LSL-*Kras*^G12D^;*Ins1*^+/−^;*Ins2*^−/−^ mice, respectively.). P vaules comparing *Ptf1a*^CreER^;LSL-*Kras*^G12D^;*Ins1*^+/+^;*Ins2*^−/−^ to *Ptf1a*^CreER^;LSL-*Kras*^G12D^;*Ins1*^+/−^;*Ins2*^−/−^ mice by two-sided unpaired t-test was 0.0306 (*). **d**, Immunohistochemical staining for CK19 in the one PDAC present in each genotype (dark blue and dark orange dots in panel c denote mice that developed tumours.). e, Relationship between composite PanIN plus tumour area and 4-h fasted insulin at 1 year of age (n= 7 for each genotype). f, The PanIN plus tumour area correlated with body weight in *Ptf1a*^CreER^; LSL-*Kras*^G12D^;*Ins1*^+/+^;*Ins2*^−/−^, but not in *Ptf1a*^CreER^;LSL-*Kras*^G12D^;*Ins1*^+/−^;*Ins2*^−/−^ mice. **g**, No correlation between PanIN plus tumour area and 4-h fasted blood glucose level at 57-weeks-old (for **f-g**, n= 13 and 9 for *Ptf1a*^CreER^;LSL-*Kras*^G12D^;*Ins1*^+/+^;*Ins2*^−/−^ mice and *Ptf1a*^CreER^;LSL-*Kras*^G12D^;*Ins1*^+/−^;*Ins2*^−/−^ mice, respectively.). **h**, Quantification of percent of total pancreatic area occupied by PanIN in male mice of each genotype at 15-30 weeks (n= 4 for each genotype). P vaules comparing *Ptf1a*^CreER^;LSL-*Kras*^G12D^;*Ins1*^+/+^;*Ins2*^−/−^ to *Ptf1a*^CreER^;LSL-*Kras*^G12D^;*Ins1*^+/−^;*Ins2*^−/−^ mice by Welch’s t-test was 0.1086. The P-value of F-test to compre variances was 0.0162(#). **i**, Relationship between PanIN area and 4-h fasted blood glucose level for male mice at 15-30 weeks (n= 4 for each genotype). **j**, Relationship between PanIN area and euthanasia age for male mice (n= 9 and 6 for *Ptf1a*^CreER^;LSL-*Kras*^G12D^;*Ins1*^+/+^;*Ins2*^−/−^ mice and *Ptf1a*^CreER^;LSL-*Kras*^G12D^;*Ins1*^+/−^;*Ins2*^−/−^ mice, respectively.). Values are shown as mean ± SEM. Sclae bars: 1 mm (**a-b** top) and 50 μm (**c** and **a-b** bottom).

We next investigated mechanisms associated with insulin-driven PanIN formation. One possible mechanism may involve the sustained mitogenic effects of insulin on PanIN cells, normal ductal cells, andlor normal acinar cells^13^. We counted the percentage of each cell type that was positive for the proliferation marker Ki67 (Fig. 3a). The variation was very high between mice within groups and there were no statistically significant differences in proliferation rate in any of the cell types (Fig. 3b). However, our data cannot exclude the possibility that insulin provided direct mitogenic and anti-apoptotic signalling to cells expressing oncogenic *Kras*^G12D^ prior to the time-points we studied.

**Fig. 3.**
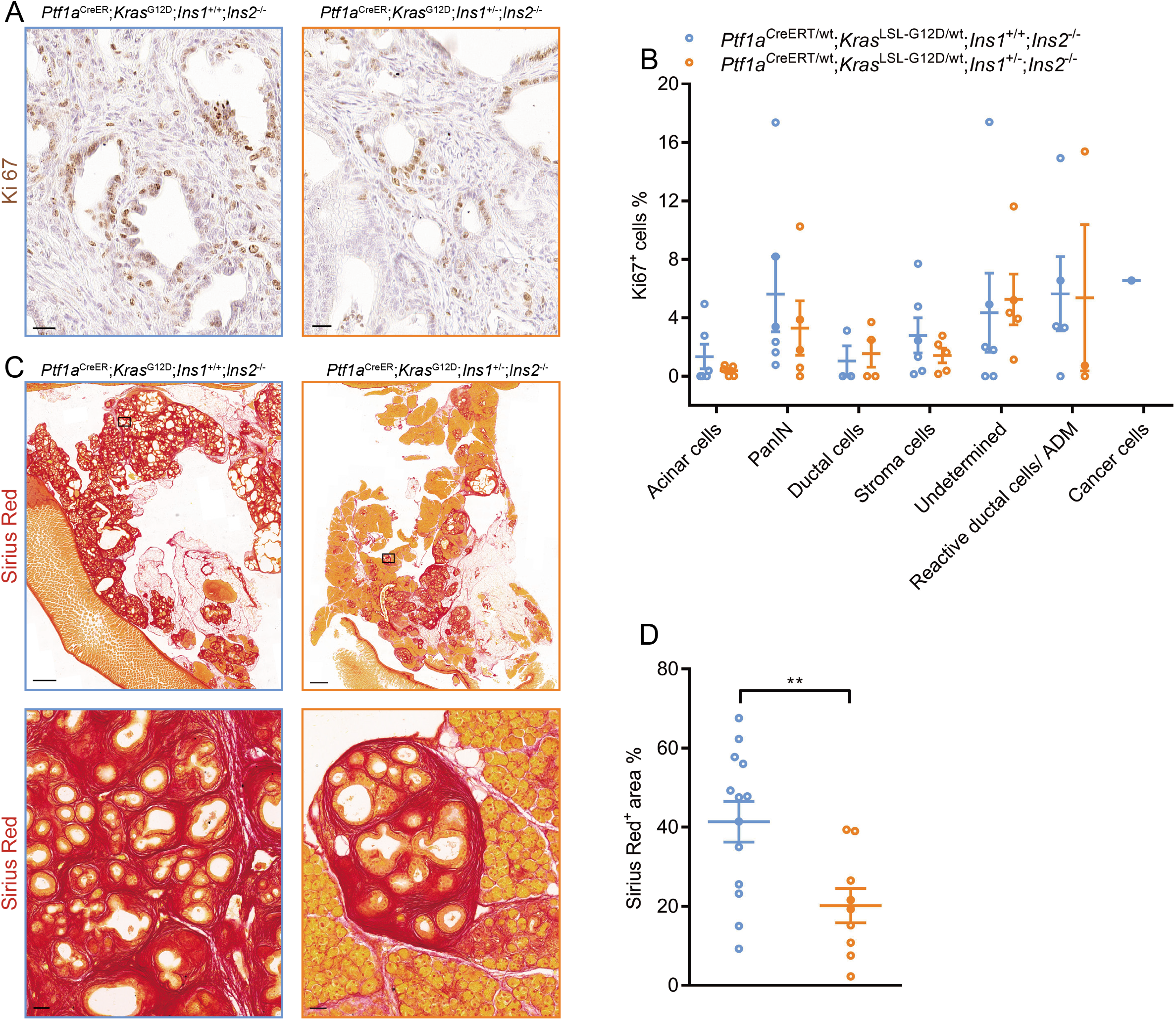
Mice with reduced insulin production have less fibrogenesis. **a**, Immunohistochemical staining for Ki67. **b**, The quantification of the percent of Ki67^+^ cells for each cell type (n= 6 and 5 for *Ptf1a*^CreER^;LSL-*Kras*^G12D^;*Ins1*^+/+^; *Ins2*^−/−^ mice and *Ptf1a*^CreER^;LSL-*Kras*^G12D^;*Ins1*^+/−^;*Ins2*^−/−^ mice, respectively.). **c**, Representative whole-section (top) and high-magnification (bottom) images of *Ptf1a*^CreER^;LSL-*Kras*^G12D^;*Ins1*^+/+^;*Ins2*^−/−^ and *Ptf1a*^CreER^;LSL-*Kras*^G12D^;*Ins1*^+/−^;*Ins2*^−/−^ pancreata stained with Sirius red. **d**, Quantification of percent of total pancreatic area occupied by Sirius Red^+^ area for each genotype (n= 13 and 9 for *Ptf1a*^CreER^;LSL-*Kras*^G12D^;*Ins1*^+/+^;*Ins2*^−/−^ mice and *Ptf1a*^CreER^;LSL-*Kras*^G12D^;*Ins1*^+/−^;*Ins2*^−/−^ mice, respectively.). P vaules comparing *Ptf1a*^CreER^;LSL-*Kras*^G12D^;*Ins1*^+/+^;*Ins2*^−/−^ to *Ptf1a*^CreER^;LSL-*Kras*^G12D^;*Ins1*^+/−^;*Ins2*^−/−^ mice by two-sided unpaired t-test was 0.0077 (**).). Values are shown as mean ± SEM. Sclae bars: 1mm (**c**, top) and 50 μm (**c**, bottom; **a**).

The second mechanism we examined was the possibility that insulin might modulate fibrosis. Insulin is known to be pro-fibrotic and previous studies have showed that insulin enhances the proliferation of pancreatic stellate cells and augments their production of extracellular matrix proteins through the PI3K pathway^21^. PDAC is characterized by an intense desmoplastic reaction resulting in highly fibrotic stroma, and this cancer-associated stroma has been suggested to support tumour survival and growth^22^. It has been proposed that HFD might exacerbate fibrogenesis and inflammation which accelerate HFD- and obesity-associated PDAC development^1, 23^. To investigate if the previously observed HFD- and obesity-induced increase in fibrogenesis were due, in part, to hyperinsulinemia, we stained for collagen with Sirius Red as a representation of fibrotic area in our pancreatic sections and quantified the percent of fibrosis area (Fig. 3c). We found that our experimental *Ptf1a*^CreER^;LSL-*Kras*^G12D^;*Ins1*^+/−^;*Ins2*^−/−^ mice had significantly less collagen deposition than control *Ptf1a*^CreER^;LSL-*Kras*^G12D^;*Ins1*^+/+^;*Ins2*^−/−^ mice (20.18± 4.34% and 41.37± 5.12%, respectively, p=0.0077) (Fig. 3d). These observations suggest that hyperinsulinemia promotes PanIN development in part by contributing to the fibrogenesis associated with PanINs.

The metabolic disturbances associated with diabetes and obesity such as hyperinsulinemia, hyperglycemia, dyslipidemia, and elevated IGF1 have been implicated, alone or in combination, in cancer risk^11, 24^. Our study is the first to separate the role of hyperinsulinemia from hyperglycemia and directly test the insulin-cancer hypothesis *in vivo*. Our data conclusively demonstrate that hyperinsulinemia, but not hyperglycemia, contributes to PanIN development through mechanisms associated with enhanced stromal fibrosis. Although hyperinsulinemia can directly affect pancreatic cells, which have robust expression of the insulin receptor, our data cannot rule out a contribution from distal tissues that may have been altered by the modest differences in circulating insulin. For example, hyperinsulinemia can promote inflammation in adipocytes and increase the levels of circulating proinflammatory cytokines like IL-6 and TNF-α^25^, potentially accelerating PanIN development. Notwithstanding, our data suggests that lifestyle interventions or therapeutics with mild insulin suppressing actions could be useful in the prevention and treatment of PDAC. The insulin lowering interventions such as exercise, low carbohydrate diets, and metformin warrant further investigation as strategies to prevent cancer or limit its progression^10, 26^.

## Methods

### Experimental Mice and Glucose Homeostasis

Animal protocols were approved by the University of British Columbia Animal Care Committee in accordance with Canadian Council for Animal Care guidelines. Mice were maintained in temperature-controlled specific pathogen-free conditions on a 12:12 hr light: dark cycle with free access to food and drinking water. *Ptf1a*^CreER^, *Kras*^LSL-G12D^, *Ins1*^−/−^, and *Ins2*^−/−^ alleles^7, 27, 28^ have previously been described. To generate background-matched experimental (*Ptf1a*^CreER^;*Kras*^LSL-G12D^;*Ins1*^+/−^;*Ins2*^−/−^ mice) and control mice (*Ptf1a*^CreER^;*Kras*^LSL-G12D^;*Ins1*^+/+^;*Ins2*^−/−^ mice), *Ptf1a*^CreER^;*Kras*^LSL-G12D^;*Ins1*^+/+^;*Ins2*^−/−^ males were bred with *Ins1*^+/−^;*Ins2*^−/−^ females. The resulting litters were fed with a high-fat diet (60% fat) (Research Diets D12492; Research Diets) at weaning (3 weeks) and were euthanized at 57 weeks. At 5-9 weeks of age, recombination was induced by four consecutive intraperitoneal injections of tamoxifen in corn oil (10mg/mL) at 0.075g tamoxifen/g body mass. Additional details of the diet composition and PCR primers used to genotype animals can be found in the Supplemental Information. Body weight, fasting glucose, and insulin were examined after 4 hours of fasting according to standard methods described previously^29^. Serum insulin levels were measured with an ELISA kit (80-INSMSU-E10; ALPCO Diagnostics, Salem, NH).

### Histological and Morphological Analysis

Pancreata were fixed for 24 hours using Z-FIX (Anatech Ltd.) followed by 48 hours with 4% paraformaldehyde and then were embedded in paraffin. Pancreata were sectioned, hematoxylin and eosin (H&E) stained, and scanned as previously described^30^. Histological analysis was conducted in a blinded fashion by Anni Zhang, Jamie Magrill, or Twan de Winter and verified by Janel Kopp and David Schaeffer (gastrointestinal pathologist). For each mouse, the PanIN area was analyzed on one of the H&E stained sections that displayed the maximal pancreatic cross-sectional area. Every gland with a lumen was categorized as normal, acinar to ductal metaplasia (ADM), PanIN, or neoplasia. Glands representing more than one of these categories were scored based on their highest-grade feature. The total pancreatic area and PanIN plus tumor area were determined by masking all pancreatic tissue and selective masking of the PanIN plus tumor area by Adobe Photoshop CC, respectively. The total pixels for pancreas or PanIN were summed using ImageJ and this was used to calculate the percentage area occupied by PanIN plus tumor per pancreas.

H&E, Alcian blue, Sirius red and immunohistochemical (IHC) staining were performed according to previously published standard methods^30^. The following primary antibodies were used: Cytokeratin 19 (rabbit, Abcam Ab133496, 1:1000) and Ki67 (rabbit, Thermo Fisher SP6, 1:200). The secondary antibody was biotin-conjugated donkey anti-rabbit (Jackson ImmunoResearch Laboratories 711-065-152, 1:500). All slides were scanned with a 20x objective using a 3DHISTECH Panoramic MIDI (Quorum Technologies Inc. Guelph, Canada) slide scanner. For Ki67 analysis, five 400 × 400 μm squares in one section per mouse was analyzed and the squares were randomly distributed in the pancreas. In total, 6 of the *Ptf1a*^CreER^;LSL-*Kras*^G12D^;*Ins1*^+/+^;*Ins2*^−/−^ mice and 5 of the *Ptf1a*^CreER^;LSL-*Kras*^G12D^;*Ins1*^+/−^;*Ins2*^−/−^ mice were randomly selected for Ki67 analysis. Anni Zhang counted the total number of cells within each square and assigned each cell into one of the following cell categories: acinar cells, PanIN cells, ductal cells, stroma cells, undetermined, reactive ductal cellsl ADM, or cancer cells. For each cell type, the number of Ki67^+^ cells and total number of cells were quantified. For Sirius red staining, one section per mouse was analyzed. The total pancreas area and fibrosis area were determined by masking all pancreatic tissue and selective masking of the Sirius red stained area by Adobe Photoshop CC, respectively. The total pixels for pancreas or fibrosis were counted by ImageJ and this was used to calculate the percentage area occupied by fibrosis per pancreas.

### Statistical Analysis

Experimenters were blinded to the genotype of mice throughout the study. All results were calculated as mean ± SEM. Statistical analyses were performed with GraphPad Prism 7.04 or the statistical programming language R (version 3.3.2). Two-tailed student’s t-test was used for normally-distributed data unless otherwise stated. Mann-Whitney test was used for the data was not normally-distributed. When comparing *Ptf1a*^CreER^;LSL-*Kras*^G12D^;*Ins1*^+/+^;*Ins2*^−/−^ and *Ptf1a*^CreER^;LSL-*Kras*^G12D^;*Ins1*^+/−^;*Ins2*^−/−^ mice’s body weight and FBG, the two-way ANOVA was used. A *p*-value < 0.05 was considered as significant.

### Data availability

All study data are available from the corresponding authors upon request.

## Acknowledgements

The authors thank members of Johnson and Kopp laboratories for discussions.

## Author contributions

AMYZ designed and managed the study, including mouse husbandry; acquired, analysed, and interpreted all data unless otherwise noted, wrote the manuscript.

JM acquired, analysed, and interpreted data (CK19, Ki67, and Alcian blue staining).

TdW acquired, analysed, and interpreted data (male PanIN staining).

XH managed the mouse colony and performed mouse husbandry.

SS provided expert advice on breeding and study design.

DFS provided expert advice on pancreatic cancer pathology and study design.

JLK analysed and interpreted the experiments, edited the manuscript, obtained funding, supervised the study.

JDJ analysed and interpreted data, edited the manuscript, obtained funding, supervised the study, conceived the study concept and design.

## Competing interests

All authors declare no competing interested related to this study.

**Correspondence and requests for materials** should be addressed to JDJ or JLK.

## Supplementary information

**Table S1.**
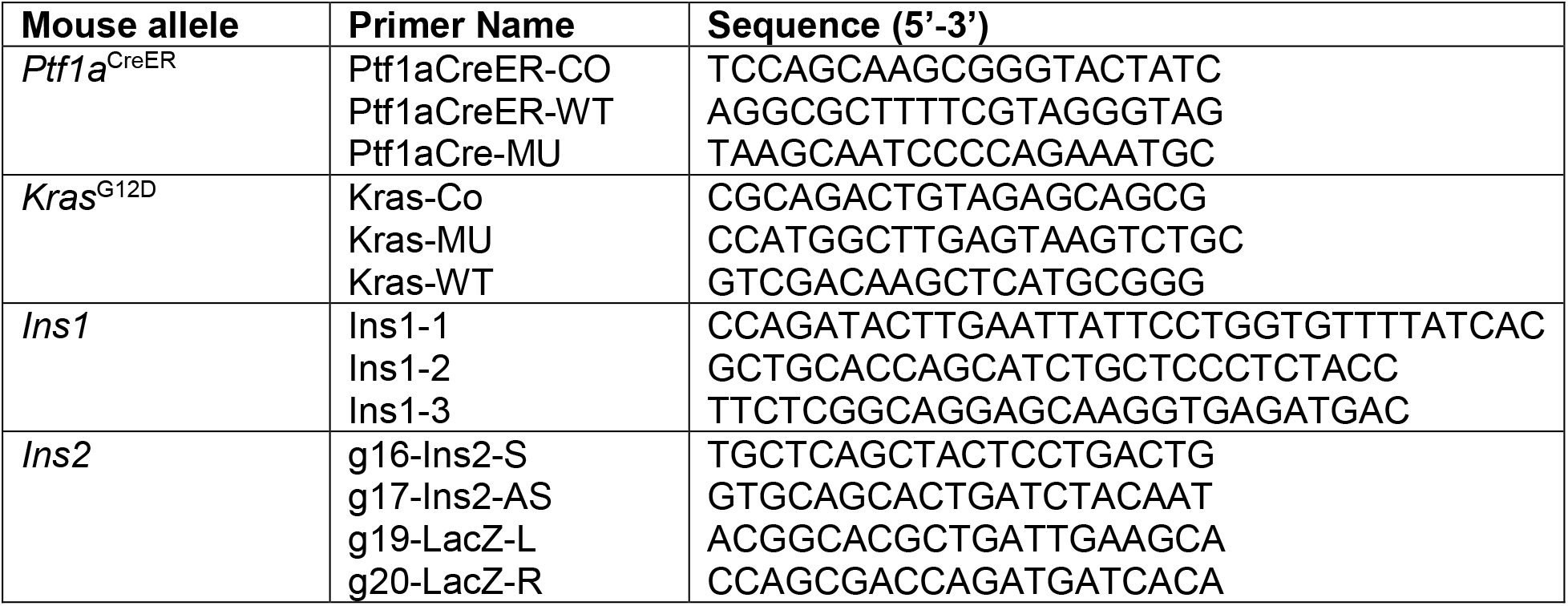
Primers used for genotyping.

